# Does retraction after misconduct have an impact on citations? A pre-post study

**DOI:** 10.1101/2020.08.11.246637

**Authors:** Cristina Candal-Pedreira, Alberto Ruano-Ravina, E Fernández, Jorge A. Ramos-Castaneda, I Campos-Varela, Mónica Pérez-Ríos

## Abstract

**Background:** Retracted articles continue to be cited after retraction, and this could have consequences for the scientific community and general population alike. This study was conducted to analyze the impact of retraction on citations received by retracted papers in two-time frames: during a post-retraction period equivalent to the time the article had been in print before retraction; and during the total post-retraction period.

**Results:** The results indicated an increase in post-retraction citations when compared with citations received pre-retraction. There were some exceptions however: first, citations received by articles published in first-quartile journals decreased immediately after retraction (*p*<0.05), only to increase again after some time had elapsed; and second, post-retraction citations decreased significantly in the case of articles that had received many citations before their retraction (*p*<0.05).

**Conclusions:** The results indicate that retraction of articles has no impact on citations in the long term, since the retracted articles continue to be cited, thus circumventing their retraction. More effective mechanisms should be established to prevent the citation of retracted articles in scientific papers.

**Methods:** Quasi-experimental, pre-post evaluation study. A total of 304 retracted original articles and literature reviews indexed in Medline fulfilled the exclusion criteria. Articles were required to have been published in Pubmed from January 2013 through December 2016 and been retracted between January 2014 and December 2016. The main outcome was the number of citations received before and after retraction. Results were broken down by journal quartile according to impact factor and the most cited papers (pre-retraction) were specifically analyzed.

## Introduction

In recent years, there has been an increase in the number of retracted articles, even when the increase in the number of indexed publications is taken into account, a development that is of concern to the scientific community (1). A paper’s retraction informs the scientific community that the study in question is not considered reliable, ethical or both, should be effectively eliminated from the literature, and should therefore cease to be cited, especially when its retraction is attributable to scientific misconduct (2). Even so, there is evidence to indicate that retracted articles continue to be cited even after their retraction (3). A classic example of this phenomenon is the case of the paper published in 1998 by Wakefield et al., which reported an association between the administration of MMR (measles, mumps, and rubella) vaccine and autism. According to a study conducted in 2019 by Suelzer et al., Wakefield’s paper had been cited a total of 1,211 times until March 2019, and of these citations, 881 occurred after the article’s retraction in March 2004 (4).

Post-retraction citations pose a problem and can have consequences, not only for the scientific community but also for the general population, since they perpetuate the misconduct and/or erroneous results, and continue lending visibility and, implicitly, credibility to the research and its authors. On the one hand, citing retracted articles may lead to other researchers accepting this information as accurate and basing their studies on it; and on the other, if health professionals make use of and base themselves on dishonest or misinterpreted information in their own clinical practice, their patients’ health could be put at risk (5). Most post-retraction citations may be attributed to two causes: first, the author’s ignorance of the retraction because, in many cases, news of the retraction does not appear in the text, or alternatively, the author does not access the article but instead places his/her trust in the citations of other authors (6,7); and second, under certain circumstances, the author may consider it useful to mention these results as erroneous in the context of his/her own study.

The frequency of retracted publications is an important indicator of both the quality of the peer review process and the pressure on researchers to publish, widely known as “publish or perish” (1,8). The number of retracted papers has risen in recent years (9)(10), as shown by the 32.2% increase in 2016 as compared to 2013 (11). The situation is rendered more serious still by the fact that only a small proportion of cases of scientific misconduct are known (12,13). For these reasons, there is a need to generate scientific evidence that would allow for systematic and rigorous analysis of the characteristics of the retracted articles and the causes of their retraction (1). A number of authors have conducted studies focusing on post-retraction citations and the impact of retraction on these, and report a reduction of 35% to 69% in the number of citations received by such articles after their retraction (5,14,15). However, these studies have neither focused on the characteristics of the journals in which the retracted articles are published, nor considered a minimum period of publication for retracted articles that would afford an opportunity for citations to be generated.

Accordingly, the main aim of this study was: to analyze whether or not citations received by retracted articles decrease after retraction, and the impact of such retraction on citations received according to the journal’s relative position (impact factor); and, to assess the impact of retraction on articles that are highly cited prior to their retraction.

## Results

A total of 304 articles fulfilled the exclusion criteria. A breakdown of the principal characteristics of the retracted articles is shown in Table 1: 65% of retracted articles were written by more than 4 authors, and 77% of these came from university institutions; China was the country of affiliation of 45.7% of first authors, followed by the USA with 14.1%; and lastly, one third of all retracted articles analyzed were published in first-quartile journals.

**Table 1.** Characteristics of retracted articles (n=304).

| Variables | n (%) |
| --- | --- |
| Country of first author |  |
| *China* | 139 (45.7%) |
| *United States of America* | 43 (14.1%) |
| *Iran* | 28 (9.2%) |
| *Other* | 94 (31%) |
| Number of authors |  |
| 1-2 | 37 (12.3%) |
| 3-4 | 69 (22.8%) |
| > 4 | 196 (64.9%) |
| Institution |  |
| *University* | 233 (77%) |
| *Hospital* | 39 (12.8%) |
| *Research centre* | 28 (9.2%) |
| *Other* | 3 (1%) |
| Scope of journal |  |
| *Biochemistry and molecular biology* | 43 (14.1%) |
| *Oncology* | 35 (11.5%) |
| *Medicine, research and experimental* | 23 (7.6%) |
| *Other* | 203 (66.8%) |
| Relative position of journal |  |
| Q1 | 100 (32.9%) |
| Q2 | 78 (25.7%) |
| Q3 | 91 (29.9%) |
| Q4 | 35 (11.5%) |
| Country of journal |  |
| *United States of America* | 97 (31.9%) |
| *United Kingdom* | 91 (29.9%) |
| *The Netherlands* | 40 (13.2%) |
| *Other* | 76 (25%) |

The citations received by retracted articles are shown in Table 2. A median of 2 pre-retraction citations were received by the articles included, with this number being equal to that received in the equivalent post-retraction period. In the case of median total post-retraction citations, however, this figure rose to 4.

**Table 2.** Characteristics of retracted articles linked to citations.

| Variables |  |
| --- | --- |
| *Median pre-retraction follow-up time and range (months)* | 18 (12-39) |
| *Median post-retraction follow-up time and range (months)* | 18 (12-39) |
| *Median total post-retraction follow-up time and range (months)* | 50 (37-72) |
| *Median total citations (P25-P75)* | 7 (3-15) |
| *Median citations in the pre-retraction period (P25-P75)* | 2 (1-5) |
| *Median citations in the post-retraction period (P25-P75)* | 2 (1-5) |
| *Median total post-retraction citations (P25-P75)* | 4 (2-8) |

Table 3 and Figure 1 shows the citations received by retracted articles according to the journal quartile in which they were published. In all cases (i.e., in both the pre- and post-retraction periods), the median number of citations was higher for first-quartile (Q1) journals. While there was a significant reduction in citations in first-quartile journals according to their BIF (*p*<0.05), this was not the case in the remaining quartiles. Median total post-retraction citations increased significantly across all quartiles, as can be seen in Figure 2.

**Table 3.** Citations received pre- and post-retraction, by journal category.

| Variables | Q1 | Q2 | Q3 | Q4 |
| --- | --- | --- | --- | --- |
| *Median total citations (P25-P75)* | 14 (7-24.25) | 6(2.75-11) | 5(2-8) | 5(2-9) |
| *Median pre-retraction citations (P25-P75)* | 5 (3-12.75) | 1 (0-4) | 1(0-3) | 1(0-3) |
| *Median post-retraction citations (P25-P75)* | 4 (2-8) | 2(1-5) | 1(0-3) | 2(1-4) |
| ***Comparison between pre- retraction period and equivalent post-retraction period*** | $p<0.001$ | $p=0.245$ | $p=0.392$ | $p=0.488$ |
| *Median total post-retraction citations (P25-P75)* | 7(4-12) | 4(1-7) | 3(2-6) | 3(1-6) |
| ***Comparison between pre- and post-retraction periods*** | $p=0.332$ | $p<0.001$ | $p<0.001$ | $p= 0.054$ |

**Figure 1.**
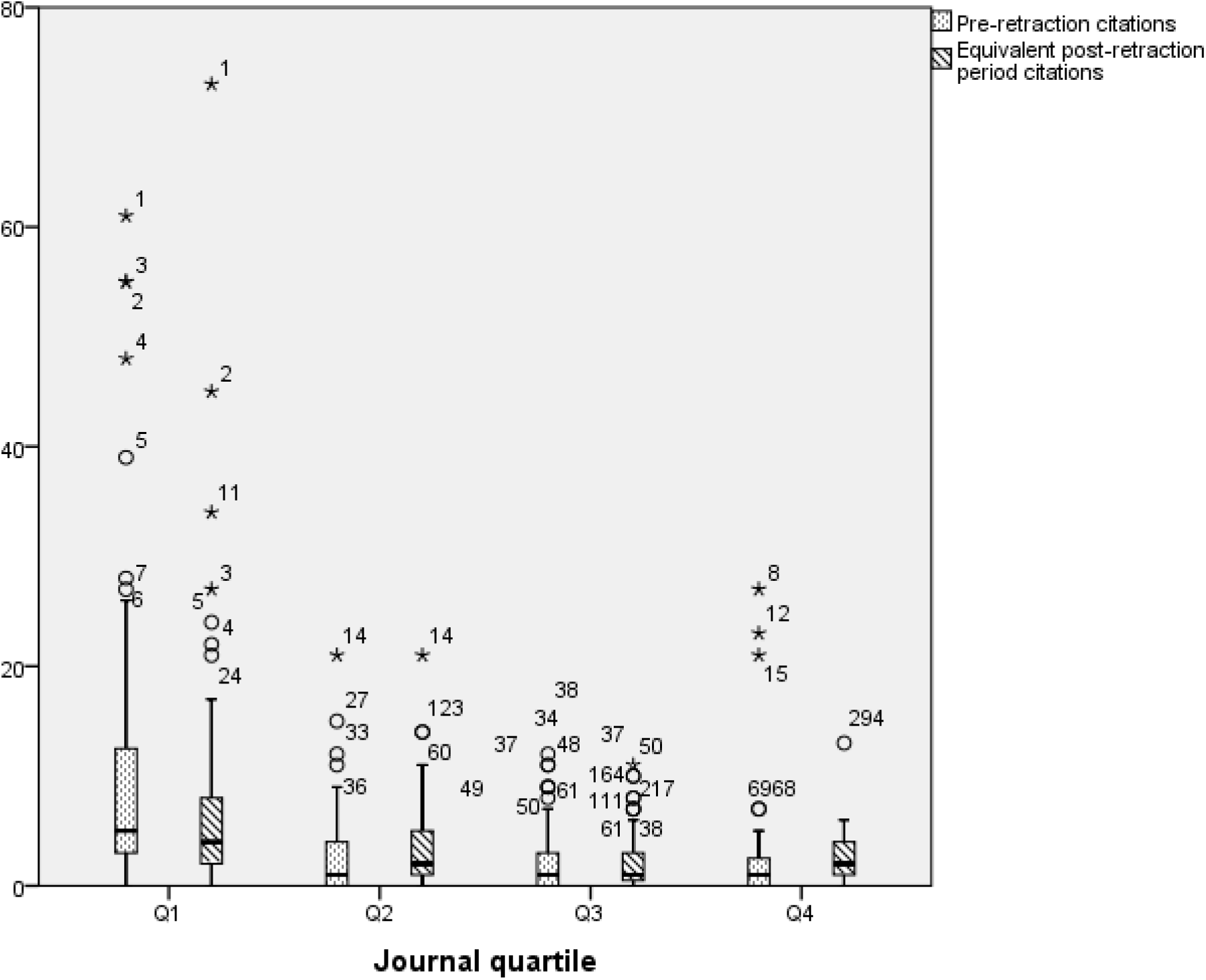
Pre- and post-retraction citations by journal quartile.

**Figure 2.**
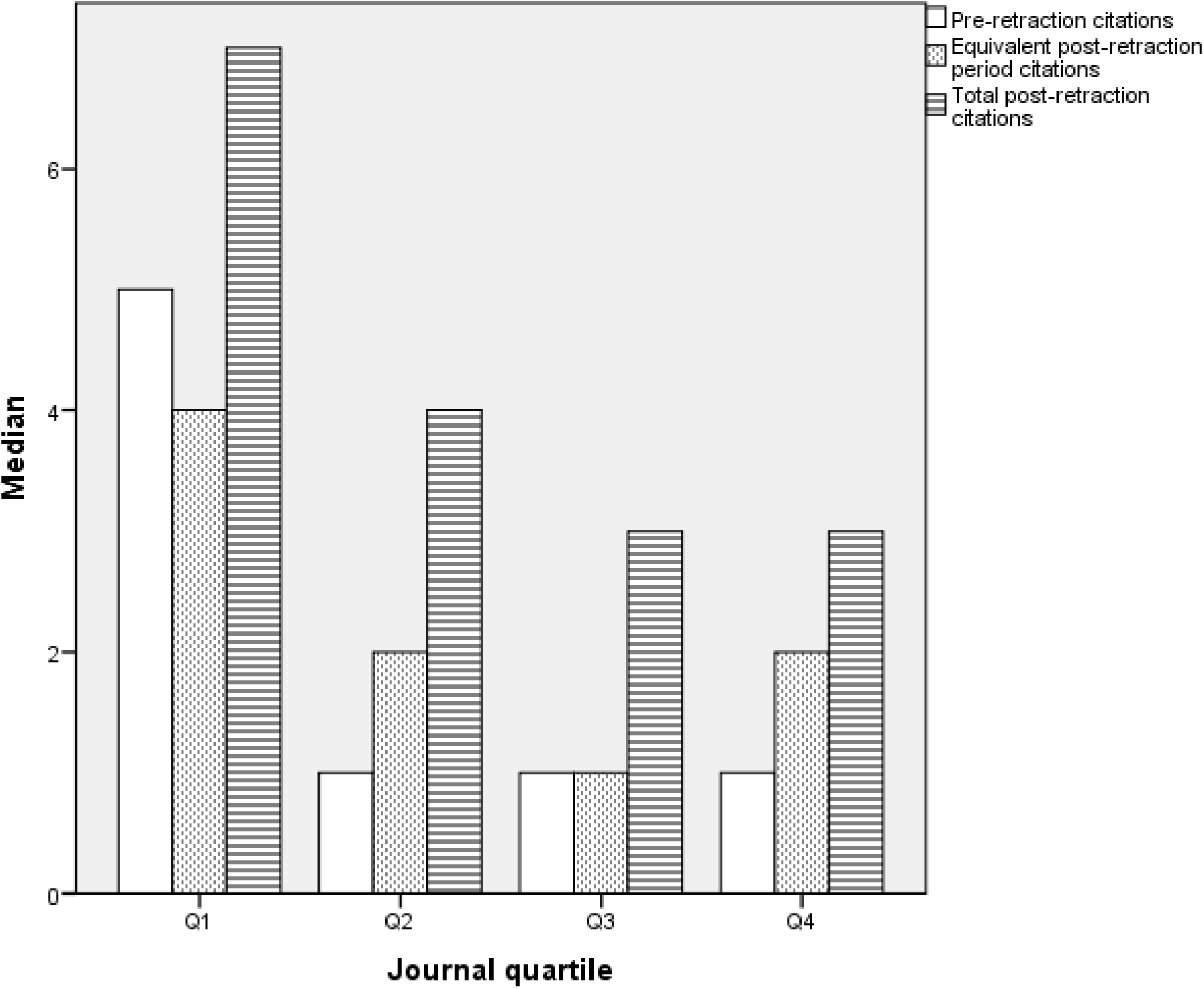
Trend in citations across the respective quartiles.

Figure 3 displays the correlation between citations received pre- and post-retraction: the scatter plot shows a positive linear correlation between the two variables, such that the more citations received pre-retraction, the more citations received post-retraction.

**Figure 3.**
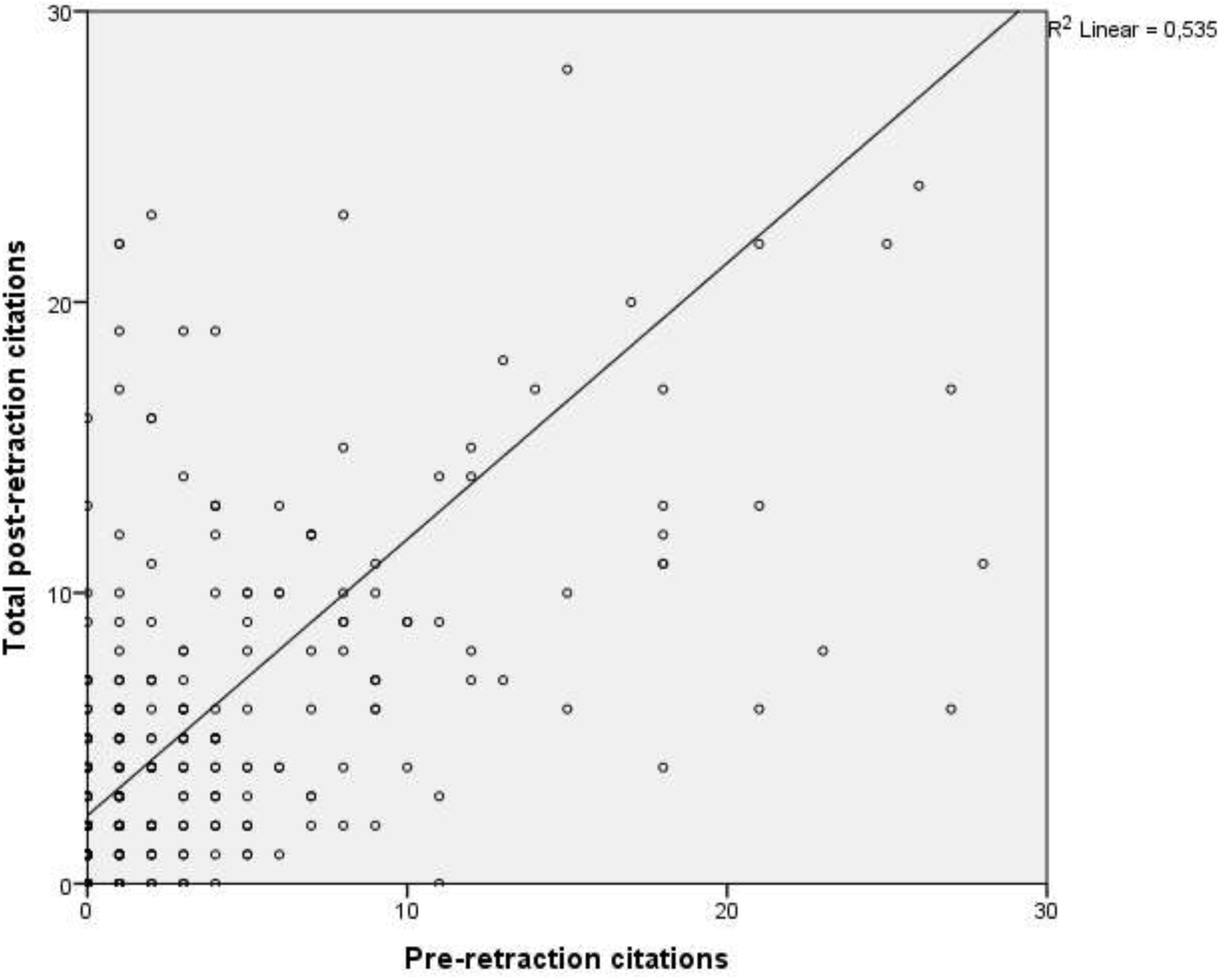
Correlation between pre- and post-retraction citations.

Table 4 describes the characteristics of the 21 most cited articles in the sample, and analyzes the citations received before and after retraction (during both the equivalent and total post-retraction periods). There was a statistically significant decrease in the median number of citations between the pre-retraction and equivalent post-retraction periods (*p*<0.05); thereafter citations registered an increase, though this was not significant (*p*=0.06).

**Table 4.** Characteristics of the most cited articles prior to retraction.

| Variables |  |
| --- | --- |
| *Median pre-retraction follow-up time and range (months)* | 27 (15-39) |
| *Median post-retraction follow-up time and range (months)* | 27 (15-39) |
| *Median total post-retraction follow-up time and range (months)* | 51 (37-59) |
| *Median total citations (P25-P75)* | 39 (30.5-70) |
| *Median citations in the pre-retraction period (P25-P75)* | 24 (18-33.5) |
| *Median citations in the post-retraction period (P25-P75)* | 13 (7.5-23) |
| **Comparison between pre-retraction period and equivalent post-retraction period** | $p=0.01$ |
| *Median total post-retraction citations (P25-P75)* | 17 (11-26.5) |
| **Comparison between pre- and post-retraction periods** | $p=0.06$ |

## Discussion

This study highlights the fact that articles retracted due to scientific misconduct continue to be cited, thus showing retraction to be an inadequate measure for eliminating invalid knowledge as a means of preventing its use by the scientific community. On the contrary, the results of this study suggest that retraction only leads to an initial decrease in the number of citations in first-quartile journals, and has no impact on articles published in second-, third- or fourth-quartile journals. To our knowledge, this is the first study with a pre-post design to be specifically targeted at analyzing post-retraction citations of individual articles retracted due to scientific misconduct, during a post-retraction period equivalent to the pre-retraction period and during the entire post-retraction period.

Different authors have studied the impact of retraction on the citations of retracted articles, with a reported reduction of 35% to 69% in the number of citations received by articles after their retraction (5,14,15). Instead of a reduction, however, the results of this study show a twofold increase in citations post-retraction. It should be stressed here that these previous studies are not comparable with ours in terms of the types of studies covered. Even so, our results coincide with those of the study undertaken by Bolboaca et al, in which most of the articles analyzed received more citations after than they did before their retraction (7).

In all, 32.3% of the retracted articles included in this study had been published in first-quartile journals. These results are in line with those of earlier studies (7,18). This may be because papers that appear in high-BIF journals and are retracted due to misconduct, are more likely to be detected, possibly owing to the additional post-publication scrutiny to which such articles are subjected as a result of being used more often than those that appear in lower-BIF journals (5,18).

With respect to citations of articles published in first-quartile journals, these decrease significantly during the equivalent post-retraction period but then subsequently increase once again, receiving more citations than those received by articles in the other quartiles. This finding is in line with those of previous studies which report a positive correlation between impact factor and post-retraction citations, namely, that the higher the journal’s impact factor, the more citations an article will receive after its retraction (15,19,20).

Different studies have found that papers which receive many citations prior to being retracted continue to be highly cited thereafter (20,20,21). This study shows a significant decrease between the citations received by the most highly cited articles prior to retraction and those received in the equivalent post-retraction period. After this period, the number of citations begins to rise again, though this increase is not significant with respect to the number received pre-retraction. However, despite the fact there is a certain impact on post-retraction citations in these types of articles, the total post-retraction citations which they receive are almost double the number of post-retraction citations received by the articles included in this study.

Bearing in mind that 94% of all post-retraction citations refer to the article as valid, without mentioning its retracted status (6), one possible explanation for the results obtained by us is that, if a retracted article is cited in a publication with no mention of its retracted status, this may give rise to “second-hand” citations. This amounts to citing an article that has previously been cited by another author, without having accessed the original and thus being ignorant of the fact that one is citing a retracted article (7). According to Simkin et al’s study, 70% to 90% of all citations are copied from references used in other articles (22). This could cause a chain reaction and thus increase citations of a retracted article long after its retraction.

One possible explanation for the decrease in citations in the immediate post-retraction period and the subsequent increase thereafter in first-quartile journals may be linked to the phenomenon of self-citations. According to the study undertaken by Madlock-Brown et al., authors may be capable of influencing the way in which their retracted paper is viewed by means of self-citation (19). Hence, there is the possibility that, in some cases, authors, on observing a decrease in citations as a consequence of retraction, cite themselves without mentioning their article’s retracted status, thereby boosting the number of citations that their retracted article receives (19).

This study has some limitations, one of which could be the low number of retracted articles. This is due to the inclusion criterion that required retracted articles to have been in print for at least one year without having been retracted. However, this inclusion criterion meant that each article had the opportunity to receive a sufficient number of citations for the impact of retraction to be measured, something that would not have been feasible if the article had been published for a short time only. Furthermore, there is the possibility that a number of post-retraction citations may have been incorrectly classified, since some came from articles accepted for publication and already available on-line before the article cited had been retracted, consequently leaving the authors unable to delete the relevant citation. Yet even if this were so, we nonetheless observed that the number of citations increased in general and that there was only an impact on journals belonging to the first quartile. This means that authors who published in high-BIF journals were aware of such retractions, in contrast to what happened in journals belonging to the other quartiles. Hence, while the period of time that elapsed between the dispatch and publication of a given article might have had some impact on the results of this study, this can be assumed to have been the same across all quartiles. An additional limitation is the fact of having only included articles published in BIF journals, which means that we do not know the results of retraction in other types of journals.

This study’s principal advantage lies in its pre-post design, since, to our knowledge, it is the first study to compare the citations received by retracted articles before retraction against those received during an equivalent period of time post-retraction. Moreover, the number of citations received, both before and after retraction, was ascertained on an article-by-article basis, thereby making it possible to ascertain the impact of different characteristics of the retracted papers on the number of citations.

In conclusion, this study indicates that retraction of articles has no impact on citations in the long term, since the retracted articles continue to be cited, thus circumventing their retraction. While it is true that, for a period of time, post-retraction citations are seen to decrease in first-quartile journals, they ultimately do start to increase again. More effective mechanisms should be established to prevent the citation of retracted articles in scientific papers: these could possibly include a warning in the publication guidelines of all scientific journals advising authors to check for citations of retracted articles.

## Data availability

The datasets generated during and/or analyzed during the current study are available from the corresponding author on reasonable request.

## Methods

### Study design

This was a quasi-experimental, pre-post evaluation study in which the intervention consisted of an article’s retraction for reasons of misconduct, and the unit of analysis was the article published and subsequently retracted. We selected original research papers and literature reviews published in journals included in Journal Citation Reports. By way of inclusion criteria, articles were required to have been published in PubMed from January 2013 through December 2016 and been retracted between January 2014 and December 2016, both inclusive. The cause of retraction must have been research misconduct, defined by the Office of Research Integrity as follows: “fabrication, falsification or plagiarism in proposing, performing or reviewing research, or in reporting research results Research misconduct does not include honest error or differences of opinion” (16).

The characteristics of retracted studies included have been described in other previous reports (11,17). All types of publications that were not original research papers were excluded, as were those retracted for any reason other than misconduct. To ensure that any given retracted article enjoyed the same opportunity of being cited pre- and post-retraction, the period during which it was in print before and after retraction was required to be equal, with a minimum pre-retraction publication period of 12 months being set as an additional inclusion criterion. This meant that all the studies included were published a minimum of 12 months prior to their retraction and were “at risk of citation” for a further 12 months thereafter, with a minimum total follow-up time of 24 months.

In view of the nature of this study, there was no need for ethics committee approval.

## Data-collection

Sociodemographic data relating to the retracted article: first author’s country of affiliation, number of authors, institutional affiliation of author(s), date of publication, and date of retraction.

Data about the journal in which the retraction occurred: category of area of knowledge (and in any case where a journal appeared in more than one such area, we selected the one in which it ranked best), bibliographic impact factor (BIF), relative position (by quartile), and country of publication.

Data on citations received: title of the citing article, name of its author(s), and name of the journal. Number of citations received by the retracted article across the total pre-retraction period, and number of citations received both in a post-retraction period equivalent to the pre-retraction period and across the total post-retraction period. In addition, we calculated total citations, defined as the sum of citations in the pre- and post-retraction periods. All citation data were obtained by conducting a manual search in the Web of Science database.

## Statistical analysis

We performed a descriptive analysis of the characteristics of the retracted articles by reference to the variables of interest, and then analyzed the association between citations received pre- and post-retraction with a breakdown by journal quartile. We first compared citations received during the pre-retraction period against those received during an equivalent post-retraction period; and we then compared these pre-retraction citations against total post-retraction citations received until December 2019. As these were matched data, the Wilcoxon signed-rank test was used for analysis purposes. Using the same method, a specific analysis was performed on the 21 most highly cited articles prior to their retraction, in order to ascertain whether their post-retraction citation pattern was affected to a relevant degree.

